# Expression patterns and interaction profiles of heterotrimeric transducin subunits in the retina of the European robin (*Erithacus rubecula*)

**DOI:** 10.64898/2026.06.29.735184

**Authors:** Srdan Vujinovic, Julia J. Forst, Sonam Kulkarni, Ümmügülsüm Güzelsoy-Flügge, Georg Langebrake, Timo Bunger, Alexander Scholten, Henrik Mouritsen, Miriam Liedvogel, Karin Dedek, Karl-Wilhelm Koch

**Affiliations:** Division of Biochemistry, Department of Neuroscience, Carl von Ossietzky Universität Oldenburg, 26111 Oldenburg, Germany; Neurosensory/Animal Navigation, Institute of Biology and Environmental Sciences, Carl von Ossietzky Universität Oldenburg, 26111 Oldenburg, Germany; Institute of Avian Research “Vogelwarte Helgoland”, An der Vogelwarte 21, 26386 Wilhelmshaven, Germany; Institute of Physics, Carl von Ossietzky Universität Oldenburg, Carl von Ossietzky Universität Oldenburg, 26111 Oldenburg, Germany; Research Center Neurosensory Science, Carl von Ossietzky Universität Oldenburg, 26111 Oldenburg, Germany

**Keywords:** Heterotrimeric GTP-binding proteins, retina, magnetoreception, RNA sequencing, immunohistochemistry, protein-protein interaction, photoreceptor cells, phototransduction

## Abstract

The heterotrimeric G-protein transducin (Gt) is mediating phototransduction in rod and cone cells of the vertebrate retina, but its expression patterns in migratory songbirds is unknown. We characterised Gt expression in the European robin, a night-migratory songbird known for its light-dependent magnetoreception. One well-supported hypothesis of magnetoreception involves radical-pair formation in the blue light receptor cryptochrome type 4a. The α- and γ-subunits of cone specific transducin have been identified as possible interaction partners of cryptochrome 4a. Therefore, we analysed the expression patterns of various G-protein subunits in bird photoreceptors by combining single cell RNA sequencing and immunohistochemistry. Protein-protein interaction was tested by pulldown, co-immunoprecipitation, and NanoBiT luminescence assays. G-protein subunits Gtβ1 and Gtβ3 are predominantly expressed in rods and cones, and GtγT2 was the principal isoform in cones, whereas Gtγ11 was associated with rods. In contrast, we did not detect Gtγ10 expression in either photoreceptor type. Interaction assays demonstrated that all three βγ combinations; βγT2, βγ10, and βγ11, can associate *in vitro*. These findings indicate that βγ dimer formation *in vivo* is likely constrained by the photoreceptor-specific expression of the respective subunits. The absence of Gtγ10 expression in rods and cones does not support a role in photoreceptor-based magnetoreception.

## 1. Introduction

European robins (*Erithacus rubecula*) are very common breeding birds which show partially migratory behaviour, with migratory individuals migrating to non-breeding areas in the Mediterranean. European robins have been demonstrated to use a light-dependent magnetic compass to orient during their migratory flights [1–3], though the sensory process of magnetoreception remains elusive. Behavioural experiments in robins show that their magnetic compass is light-dependent [1,3], that this compass can be disrupted by radio-frequency fields predicted to affect a radical-pair-based mechanism [4–8], and the eye is likely involved in the reception process[1,9,10]. In the bird’s outer retina, light is transduced by two cell classes, rods and cone photoreceptors. Bird photoreceptors comprise four types of single cones [11] sensitive to different wavelengths of light (red, green, blue and UV, recently termed P1-P4, respectively, [12], a double cone (consisting of a principal and an accessory member, P5 and P6, respectively), and a type of rod (P0). Double cone photoreceptors have recently been proposed as a potential location for magnetoception [13–15], with the accessory member of the double cone as the most likely primary sensory cell [15]. European robin double cones are organised in a highly ordered mosaic, suggested to facilitate the separation of magnetic directional information from the Earth’s magnetic field from differences in intensity and light polarization [13,16]. Schulten et al. (1978) [17] proposed a light-triggered magnetosensitive radical-pair mechanism as the molecular basis of magnetic field detection.

Later, Ritz et al. (2000) [5] suggested cryptochrome proteins as candidates matching the required characteristics of a blue-light photoreceptor. Cryptochrome 4a (Cry4a) is currently the best supported magnetoreceptor candidate in the European robin. Purified recombinant *Er*Cry4a binds the light-absorbing co-factor Flavine Adenine Nucleotide (FAD) and is magnetically sensitive *in vitro* [18]. Moreover, *Er*Cry4a is expressed in red cones and double cones of the European robin retina [14], suggesting that the primary steps of magnetoreception are initiated alongside the phototransduction cascade.

Several Cry4a interacting proteins, which could potentially be part of a downstream signalling cascade, have been identified by yeast-two-hybrid screening [19]. Among those identified candidates were the genes *GNAT2* and *GNG10* coding for the α-and γ-subunit of a heterotrimeric G protein, respectively. Out of all isoforms of α-subunits, the same Gtα-subunit encoded by *GNAT2* has been detected across all cone types in the chicken retina [20], where it plays a central role in the phototransduction cascade [21–23]. In rods, *GNAT1* regulates the expression of rod-specific Gtα. However, there are at least 5 types of β- and 12 types of γ-subunits across different vertebrate species [24], yet *GNB1* encodes the rod β-subunit, and *GNB3* encodes the cone β-subunit across vertebrates [25]. The larger variety of γ-subunits implicates a complex diversity of subunit combinations present in different signalling pathways. Several γ-subunits are connected to phototransduction, as *GNGT1* encodes a rod-specific and *GNGT2* a cone specific subunit in several species [26,27]. Other studies located the *GNGT2* gene in mammals in tandem with *GNGT1*, suggesting a local duplication of *GNGT1* in the mammalian lineage [28]. In contrast, *GNG11* seems to be encoding a rod γ-subunit [26,27]. *GNG11* is present in all placental mammals, but was not identified in some marsupials and possibly arose in a mammalian common ancestor [26]. It is apparently missing in other vertebrate groups, including birds. Both *GNGT1* and *GNGT2* genes were identified in genomes of all major vertebrate taxa, including bony fish, amphibians, birds and reptiles [26], and a recent transcriptome analysis also reveals expression of *GNGT1/GNG11* in chicken rods [29,30]. Along with these results, a strong expression of the *GNGT2* transcript was also reported for both rods and cones of chicken [29,30]. Therefore, both *GNGT1/GNG11* and *GNGT2* subunits are co-expressed in rods of the chicken retina, while *GNGT2* is only expressed in cones. However, nothing is known about the γ-subunit expression in songbirds.

Wu et al. (2020) [19] identified *GNG10* (γ10) as a putative Cry4a-interacting protein, but its presence and location in the bird retina remained unknown. Expression analyses revealed that γ10 is not primarily expressed in mammalian rod or cone photoreceptor cells [31,32]. According to these studies, the cone specific γ-subunit is encoded by the gene *GNGT2*. Strong interaction of the cone specific β-subunit β3 (gene *GNB3*) was observed for γ11, but not for γ10 [33], which makes combinations of a βγ-complex in the European robin retina distinct from β3γ10 likely.

The hypothesis that two different senses, vision and magnetoreception, could operate in photoreceptor cells to guide migration in songbirds requires the investigation of the expression and localisation patterns of the potentially involved signalling molecules on the cellular level. Previous studies on the expression of heterotrimer G protein subunits were not carried out in species for which magnetic sensing capabilities have been documented. For migratory songbirds like European robins, it is therefore not known which G protein subunits, or which combinations of subunits are present in which photoreceptor cell types. Here we address these questions by a combination of transcriptome cluster analysis, immunohistochemistry, co-immunoprecipitation, and a luminescence-based protein-protein interaction assay. This work presents, to the best of our knowledge, the first study of G-protein subunit expression in migratory songbirds, aimed to give a deeper understanding of G-protein variety in the bird retina as well as G-protein subunit complex formation relevant for vision and putatively also for magnetoreception.

## 2. Results

### 2.1. Expression of G Protein subunit transcripts in European robin photoreceptors

To analyse the G-protein subunit transcripts in photoreceptors, we performed RNAseq on samples from the European robin retina. Cell types were identified using Seurat-based clustering and annotation, showing distinct transcriptional populations (Figure S1, Table S1). For this analysis, we classified the cells into the broad cell types of photoreceptors (rods and cones), ON and OFF bipolar cells, horizontal cells, amacrine cells, and retinal ganglion cells, as well as oligodendrocytes (not shown) and Müller glia cells (Figure 1).

**Figure 1.**
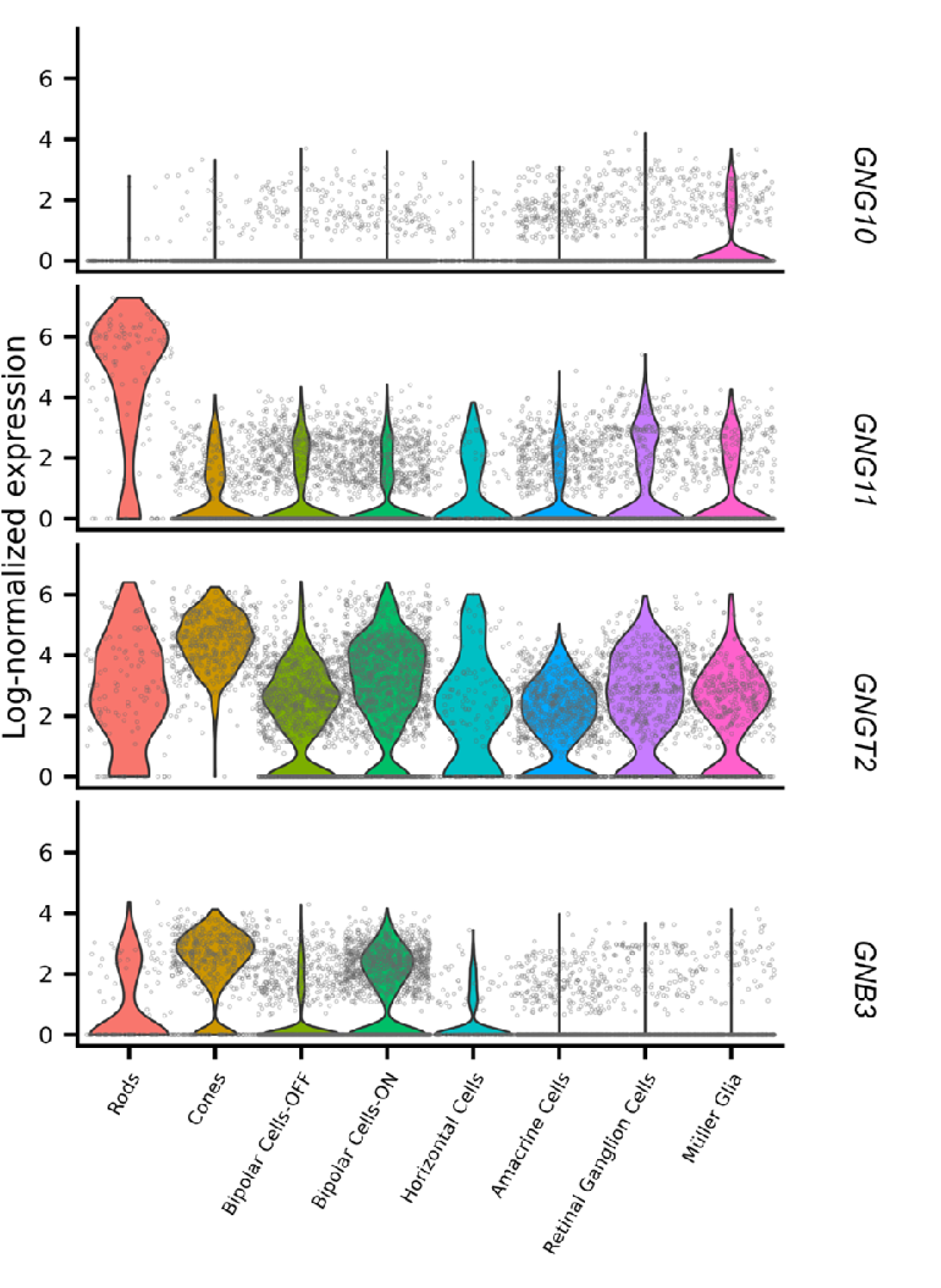
Violin plots showing the expression levels of *GNB3, GNG10, GNG11* and *GNGT2* in different retinal cell types. Expression levels are plotted as log-normalized transcript counts.

Expression analysis of each G-protein subunit revealed that *GNB1* and *GNG11* show high levels of expression in the rod photoreceptors. *GNB3* is expressed more in the cone photoreceptors, while *GNB5 is* expressed in rods and cones, with a higher fraction of cones expressing the subunits. All the cell types investigated expressed *GNGT2*, with the highest expression in cones. The remaining subunits investigated, i.e. *GNB4, GNG2, GNG7, GNG10* and *GNG12,* were not significantly expressed in photoreceptor cells (Figure 1, Supplementary Figure S2, S3, Table 1)

**Table 1.** The mean log-normalized counts of each gene per cell i.e. the expression level, Log2 Fold Change and the Benjamini-Hochberg adjusted p-values for the expression of each G-protein subunit in the two photoreceptor classes-the rods and cones.

|  | Gene | Mean log normalized expression | Mean log2Fold Change | Adjusted p-value |
| --- | --- | --- | --- | --- |
| Rods | *GNB1* | 2.797368351 | 3.542842 | 1.79E-43 |
|  | *GNB3* | 0.919537584 | 0.1033851 | 1.00E+00 |
|  | *GNB4* | 0.062830394 | -2.1533189 | 1.00E+00 |
|  | *GNB5* | 1.321833042 | 0.7676815 | 1.15E-06 |
|  | *GNG2* | 0.091318065 | -1.8279051 | 1.00E+00 |
|  | *GNG4* | 0.085091153 | -1.2267109 | 1.00E+00 |
|  | *GNG7* | 0.245788848 | -0.5303993 | 1.00E+00 |
|  | *GNG10* | 0.060921243 | -1.8985638 | 1.00E+00 |
|  | *GNG11* | 4.487924778 | 5.8503137 | 1.24E-48 |
|  | *GNG12* | 0.035413621 | -0.4533134 | 1.00E+00 |
|  | *GNGT2* | 2.808915897 | 0.4393597 | 1.00E+00 |
| Cones | *GNB1* | 0.246267831 | -1.8623156 | 6.92E-23 |
|  | *GNB3* | 2.32107783 | 2.0335096 | 8.78E-160 |
|  | *GNB4* | 0.171979476 | -0.1484593 | 1.00E+00 |
|  | *GNB5* | 1.812445584 | 1.9174565 | 5.06E-137 |
|  | *GNG2* | 0.070622496 | -2.3436673 | 4.81E-11 |
|  | *GNG4* | 0.045746025 | -1.9001579 | 2.20E-03 |
|  | *GNG7* | 0.260516343 | -0.3955034 | 1.00E+00 |
|  | *GNG10* | 0.103211169 | -1.1683369 | 3.65E-02 |
|  | *GNG11* | 0.829866109 | -1.6110642 | 1.00E+00 |
|  | *GNG12* | 0.011044816 | -1.7027505 | 1.00E+00 |
|  | *GNGT2* | 4.447627316 | 1.9491321 | 5.13E-169 |

### 2.2. Expression of G protein subunit proteins in European robin photoreceptors

Earlier studies [19,34] suggested Gtα2 as an interaction partner of the putative magnetosensory protein cryptochrome 4a (Cry4a). If Gtα2 and Cry4a interact in retinal tissue, they need to be expressed in the same cell type and subcellular compartment. Indeed, double labelling with antibodies for Gtα2 and Cry4 showed that both proteins colocalise in the outer segments of photoreceptors (Figure 2), which seem to belong to red cones and double cones [14]. Please note that anti-Cry4 antibodies cannot differentiate between the two known Cry4 isoforms [14], Cry4a and Cry4b [35], which is why we refer to Cry4 in antibody stainings. However, the staining almost certainly detects Cry4a, since a parallel study suggested that Cry4b is unlikely to be translated into a protein in the European robin [36]. Antibodies for Gtα2 additionally labelled the inner segments and the end feet of photoreceptors (Figure 2, arrows) whereas Cry4 labelling was largely restricted to the outer segments.

**Figure 2.**
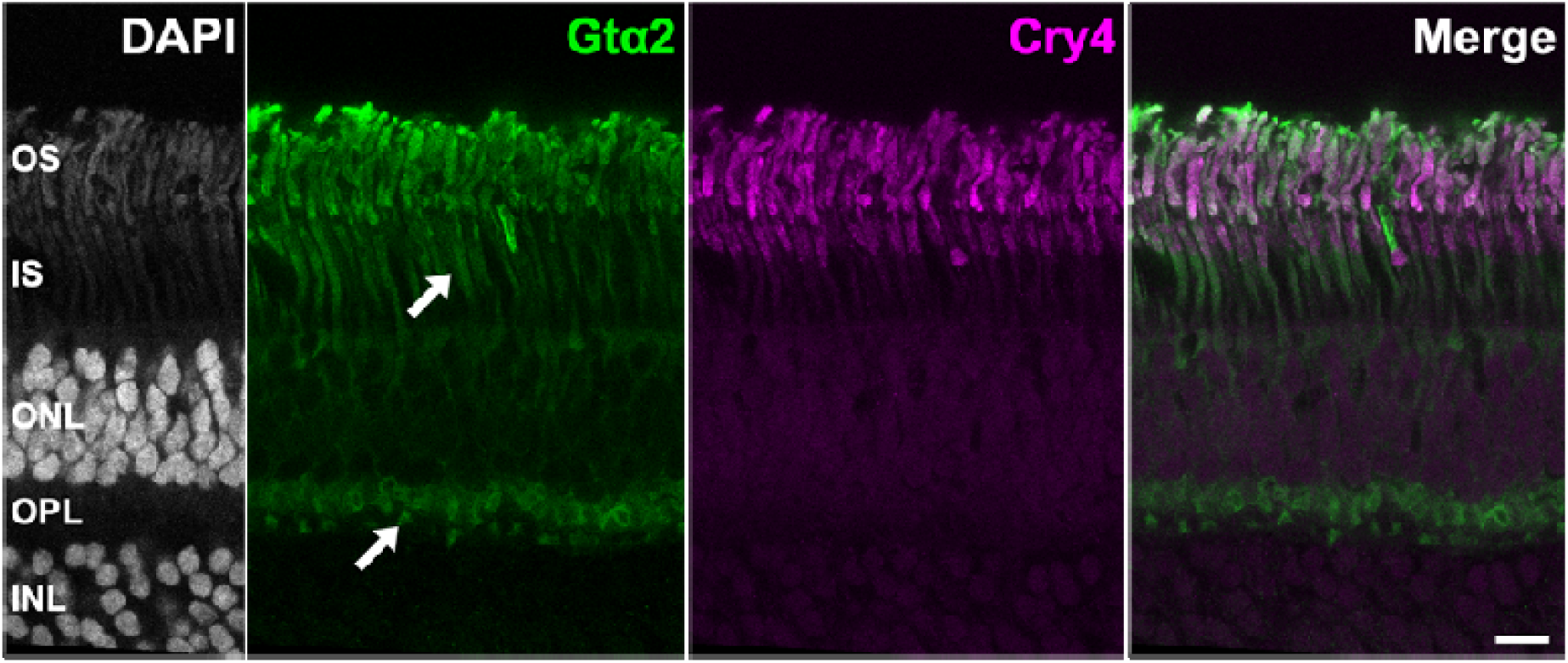
Immunohistochemical double labelling of European robin retinal sections for Gtα2 and Cry4. Gtα2 and Cry4 immunoreactivity colocalized in the outer segments of photoreceptors, likely of red cones and double cones. In addition to outer segments, Gtα2 was also detected in the inner segments and photoreceptor endfeet (arrows). Cry4 labelling was largely restricted to the outer segments. DAPI labeled sections (gray-scale) to visualize cell nuclei and retinal layers. OS, outer segments; IS, inner segments; ONL, outer nuclear layer; OPL, outer plexiform layer; INL, inner nuclear layer; Scale bar: 10 µm.

The β-subunit interacting with Gtα2 in the bird retina is most likely Gtβ3, as it was shown to be expressed in all photoreceptors and ON bipolar cells in the chicken retina [25,37]. Here, we confirmed photoreceptor expression in the European robin retina as well. Gtβ3 labelling was present in all photoreceptor outer segments and was particularly strong in some photoreceptors (Figure 3, arrows). These strongly Gtβ3-positive photoreceptors did not express Cry4 and consequently, do not represent red or double cones [14]. Based on their morphology, these cells are most likely rods.

**Figure 3.**
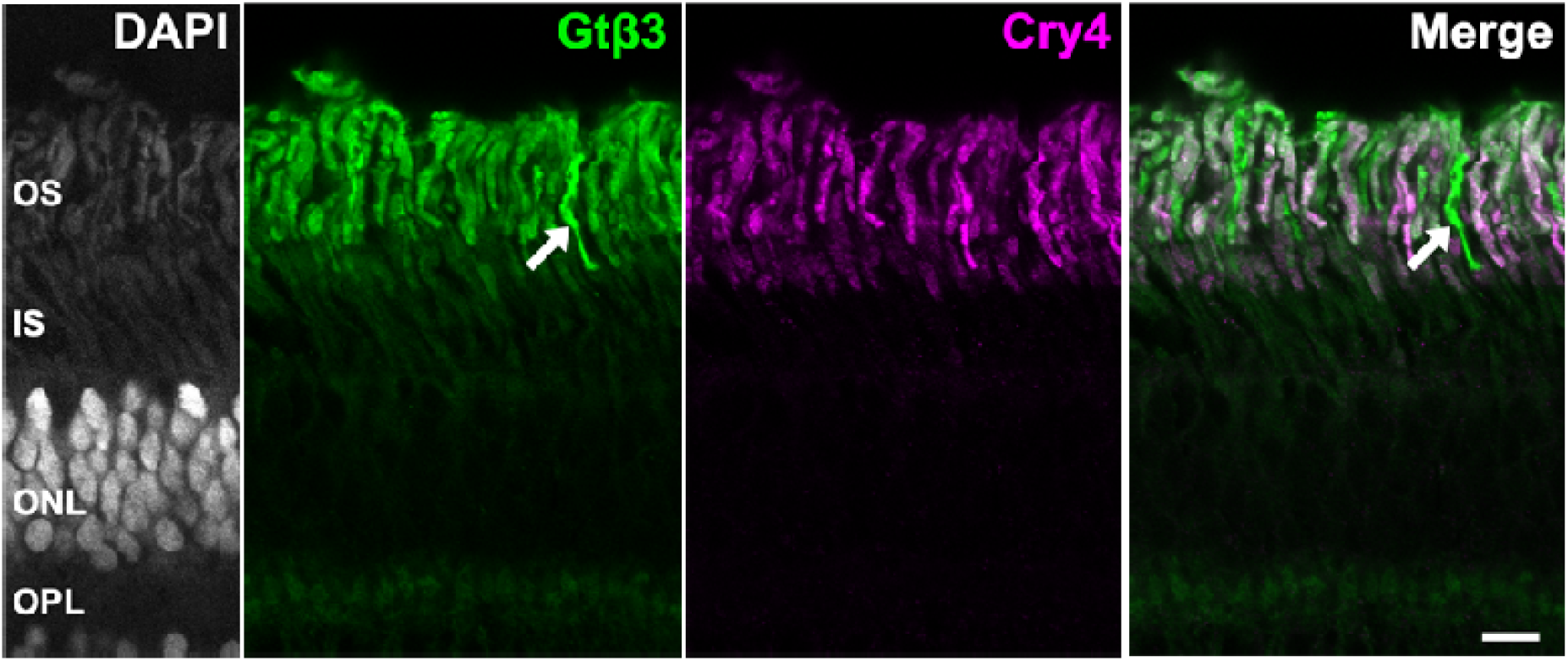
Immunohistochemical double labelling of central European robin retina for Gtβ3 paired with Cry4. Gtβ3 labelling was present in all photoreceptor outer segments, with particularly strong labeling in a subset of cells (arrows). These strongly Gtβ3-positive photoreceptors lacked Cry4 immunoreactivity, indicating that they are neither red cones nor double cones. DAPI labeled sections (gray-scale) to visualize cell nuclei and retinal layers. OS, outer segments; IS, inner segments; ONL, outer nuclear layer; OPL, outer plexiform layer; Scale bar: 10 µm.

To analyse the expression of G protein γ-subunits at the protein level, we stained slices of the European robin retina with antibodies directed against GtγT2, Gtγ11, and Gtγ10, which we identified by scRNAseq as being expressed in rods and cones. In line with the transcriptomics data, no specific signal was detected with the Gtγ10 antibody (Supplementary Figure S4). The antibody for GtγT2 revealed strong signals in the outer segments of photoreceptors, whereas inner segments and synaptic pedicles showed faint GtγT2 signals (Figure 4A, arrows). GtγT2 labelling colocalized with Gtβ3 immunoreactivity (Figure 3) and with labelling for rhodopsin and long-wavelength opsin (Figure 4A), indicating expression in rods as well as in red cones and double cones [11], respectively.

**Figure 4.**
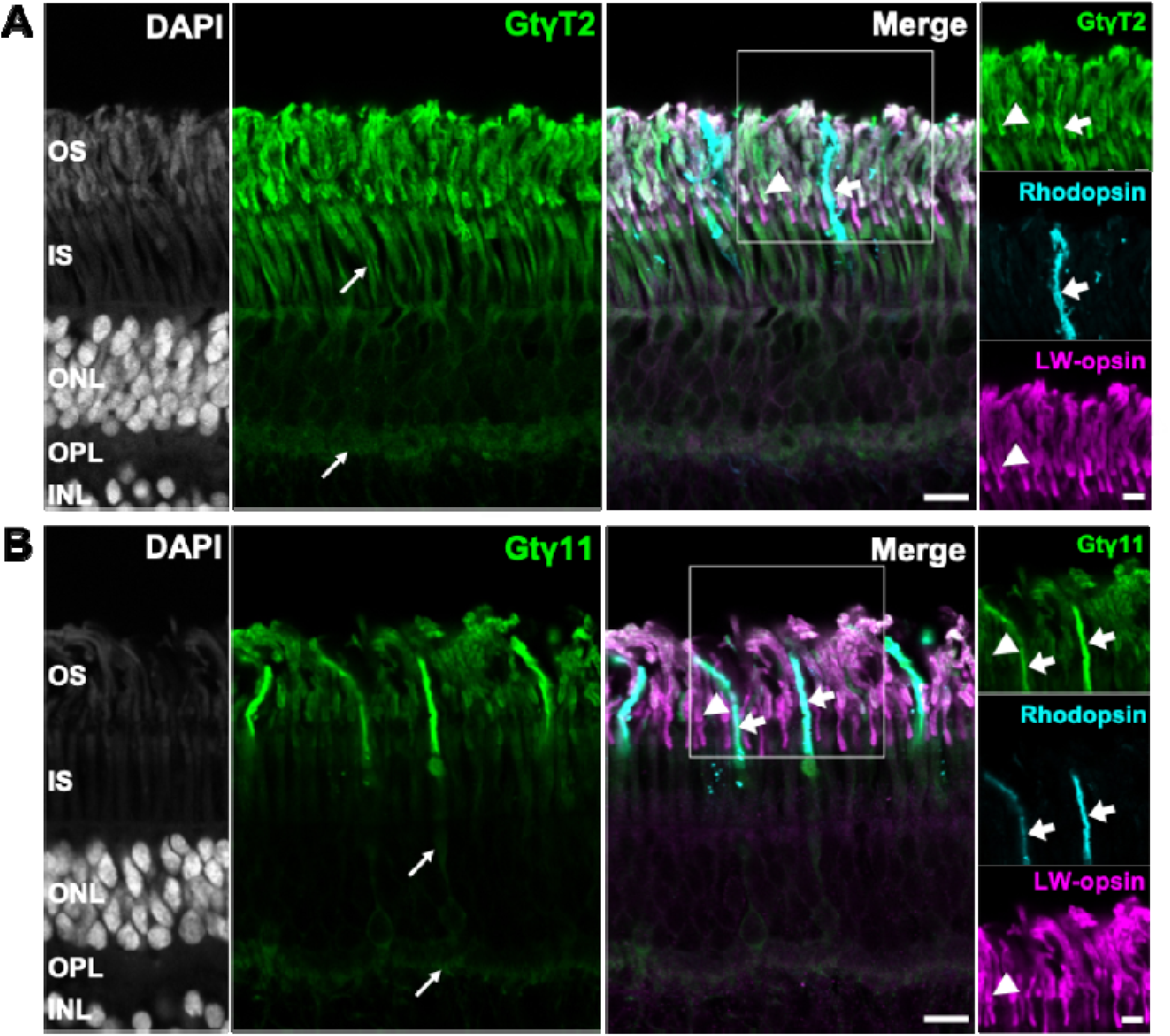
Immunohistochemical triple labelling of central European robin retina for Gtγ-subunits GtγT2 (A) and Gtγ11 (B), each paired with Rhodopsin and LW-opsin. Boxed regions are shown as close-up channel overlays. The GtγT2 immunoreactivity was strongest in photoreceptor outer segments, where it overlapped with Rhodopsin and LW-opsin. Weaker labelling was also detected in inner segments and photoreceptor endfeet (A, arrows). Gtγ11 strongly labelled rods, as demonstrated by the complete overlap with Rhodopsin-positive cells (B, arrows), whereas weaker immunoreactivity was observed in red cones and double cones, indicated by the complete overlap with LW-opsin-positive cells (B, arrowhead). DAPI labelled sections (gray-scale) to visualize cell nuclei and retinal layers. OS, outer segments; IS, inner segments; ONL, outer nuclear layer; OPL, outer plexiform layer; INL, inner nuclear layer. Scale bars: 10 µm. Close-ups: 5 µm.

Antibodies for Gtγ11 revealed strong staining in the outer segments of rod photoreceptors (Figure 4B, arrows) as evidenced by the complete overlap with rhodopsin labelling. The weaker Gtγ11 staining (Figure 4B, arrowheads) colocalized with LW-opsin, suggesting a weak expression in red cones and double cones [11].

In summary, we revealed a colocalization of Cry4 and its putative interaction partner Gtα2, confirmed Gtβ3 expression also for the European robin retina, detected GtγT2 weakly and Gtγ11 strongly expressed in rods, and showed GtγT2 to be expressed in both rods and cones. These results largely confirm the scRNAseq data.

### 2.3. Pulldown and Co-immunoprecipitation

The subunit composition of the heterotrimeric G-proteins in photoreceptors could either be determined by a differential expression pattern or by subunit-specific interactions. To test for this, we used *ExpiSf9 cells* to coexpress Gtβ3 and the three Gtγ-isoforms to test which of these proteins can form a heterodimer. Cell pellet lysates containing the Gtβ3-subunit and His-tagged Gtγ-isoforms were applied to a Ni^2+^-chelating column (Figure 5A). Bound protein complexes that were caught on the Ni^2+^-column were eluted with a buffer containing 300 mM imidazole. Eluted samples were analyzed by SDS-PAGE (Figure 5A). The size of the Gtβ-subunits on the gel is ∼35 kDa, which agrees with theoretical mass, considering that the samples are run in 15% SDS-gel. Gamma-subunits are present around 12 kDa, which corresponds to theoretical mass of His-tagged Gtγ-subunits (7.7 kDa for Gtγ10, 8.7 kDa for Gtγ11, 8.3 kDa for GtγT2, Supplementary Table S2). Differences are observable for Gtγ10 compared to Gtγ11 and GtγT2, due to smaller sizes. Beta-subunit 3 is not present in non-transfected cells or in cells expressing only Gtβ3 as well as γ-isoform (lane β3 in Figure 5A), which confirms that the heterodimer binds only to the column when it is coexpressed with a His-tagged Gtγ-isoform. Additional bands in the SDS-PAGE analysis represent background signals due to unspecific binding of other cellular proteins to the Ni^2+^-resin. By using Protein G Dynabeads with covalently bound anti-His antibody, we observed less nonspecific binding of cellular proteins to the beads and obtained a specific elution of heterodimers (Figure 5B). Immunoblotting confirmed the presence of Gtβγ-dimers in eluates by using specific anti-Gtβ and anti-Gtγ antibodies and the corresponding secondary antibodies (Figure 5B). No detection of Gtβ3 was seen in the eluates in the absence of Gtγ-subunits, independent of whether we employed the Ni^2+^-resin (Figure 5A) or the Protein G Dynabeads (Figure 5B). The results showed that Gtβ3 can form a heterodimer complex with all three Gtγ-subunits. Interestingly, the interaction was possible without the CAAX motif in the C-termini of the Gtγ-subunits necessary for prenylation. Presence of the prenyl-group in the Gtγ-subunits, however, did not change the results (see below under NanoBiT-assay).

**Figure 5.**
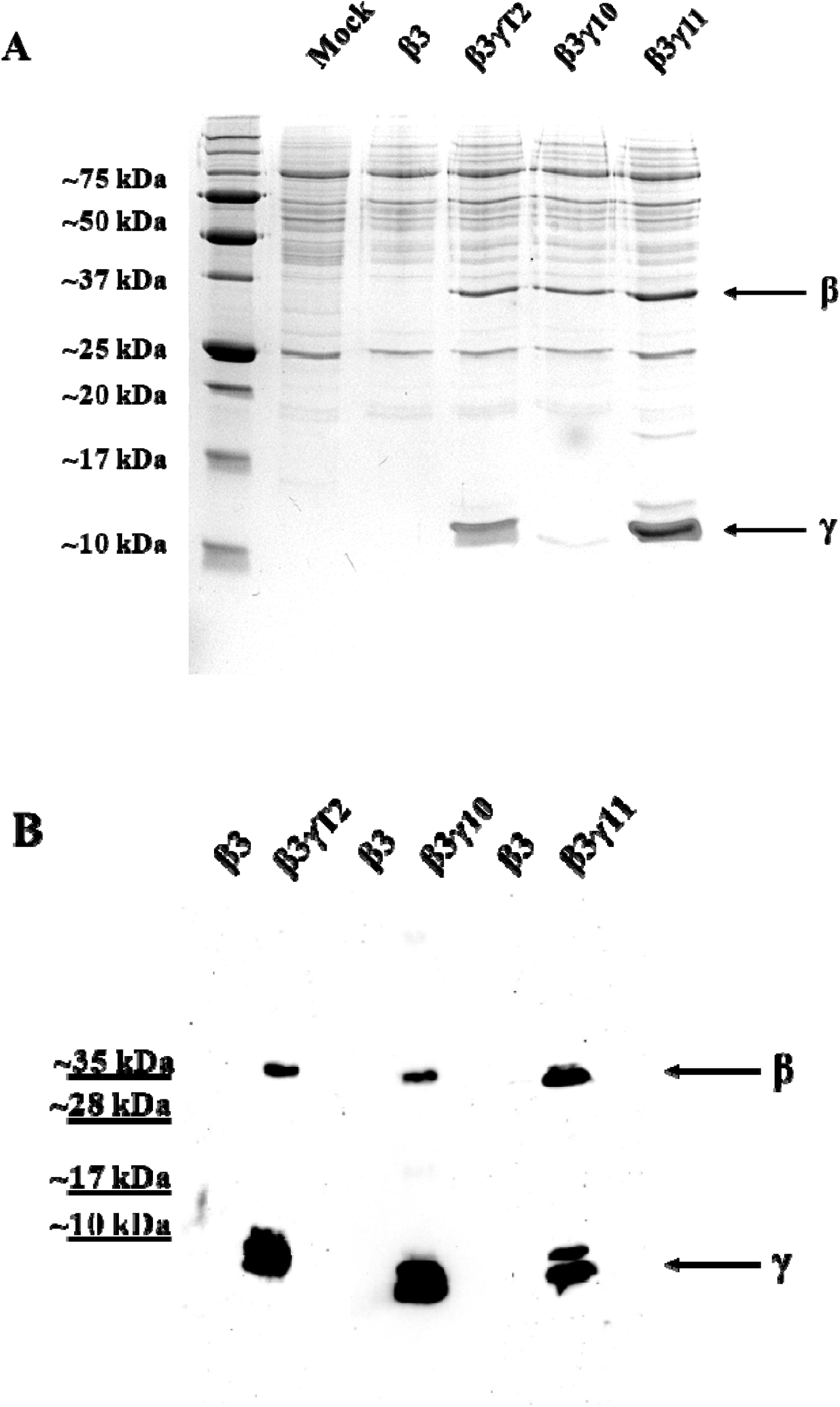
A: Pulldown of Gtβγ subunits with Ni^2+^-resin to test for Gtβγ-subunit interaction in solution. SDS-PAGE was performed on a 15% SDS gel followed by Coomassie staining. As negative controls, non-transfected ExpiSf9 cells and only Gtβ3-subunit expressing cells were used. The γ-subunits contained a 6-His Tag that allows pulldown with Ni-resin. The β-subunit 3 could be detected in all 3 samples when it was coexpressed with Gtγ-subunits. B: Western Blot of eluates from coimmunoprecipitation using His-antibody-coupled Dynabeads. Antibodies directed against Gtβ3, GtγT2, and Gtγ10 have been used to detect the subunits (Table 1). For the Gtγ11 subunit, an anti-His antibody was used, since the antibody directed against Gtγ11 produced weaker signals. In samples expressing only the Gtβ3 subunit, the Gtβ3 subunit could not be detected. Detection was observed only when the Gtβ3 subunit was co-expressed with Gtγ subunits.

### 2.4. NanoBiT Assay

To validate the results from the pull-downs and co-immunoprecipitation, an additional method was employed to test heterodimer formation of Gtβ3 with different Gtγ subunits. The luminescence based NanoBiT-assay is a protein-fragment complementation assay and allows protein-protein interactions in living cells (e.g. HEK293 cells). A further advantage of the assay is the combination of high sensitivity and low background. First, we tested the expression of the constructs by Western blotting using an anti-LB antibody (see Table 2 and sections 4.4 and 4.7 in the Methods part), allowing detection of each LB-fused subunit (see Methods). In Figure 6A, LB-Gtγ constructs are visible at ∼25.4 kDa (18.1 kDa for LB and 7.3 kDa for Gtγ subunit). For the LB-Gtβ construct, the size was ∼55.3 kDa (18.1 kDa for LB and 37.2 for Gtβ-subunit). The results of the NanoBiT assay are presented in Figure 6B. An increase in luminescence signals was measured for all three tested pairs: Gtβ3γT2 (orange bars), Gtβ3γ10 (yellow bars) and Gtβ3γ11 (green bars) when compared to the negative control (HaloTag constructs). All signals of the analyzed SB-Gtβ:LB-Gtγ pairs were about 100 times higher in relative luminescence than the negative control samples that lacked the Gtβ-subunit part. Some pairs showed even higher luminescence than the positive control (blue bar: LB-PKA RIIα:SB-PKA Cα). Switching the localization of the NanoLuciferase tags in the constructs resulted in LB-Gtβ:SB-Gtγ pairs with lower amplitudes of luminescence signals (columns on the right side in Figure 6B). This could potentially point to a less efficient reconstitution of NanoLuciferase. However, these relative signals were also nearly 100 times higher than the control construct. Only the Gtγ10 construct showed a lower, ∼10-fold increase. The complementary fragments of NanoLuciferase (LB and SB) associate with very low affinity (K_D_ around 190 µM), so spontaneous reconstitution is not likely to occur unless the tested proteins are in close proximity. Thus, our measurements showed a high signal-to-noise ratio supporting our pulldown and co-immunoprecipitation experiments (Figures 6A and 6B). The results further demonstrated that Gtβγ-dimer formation occurred in living HEK293 cells and could be measured.

**Figure 6.**
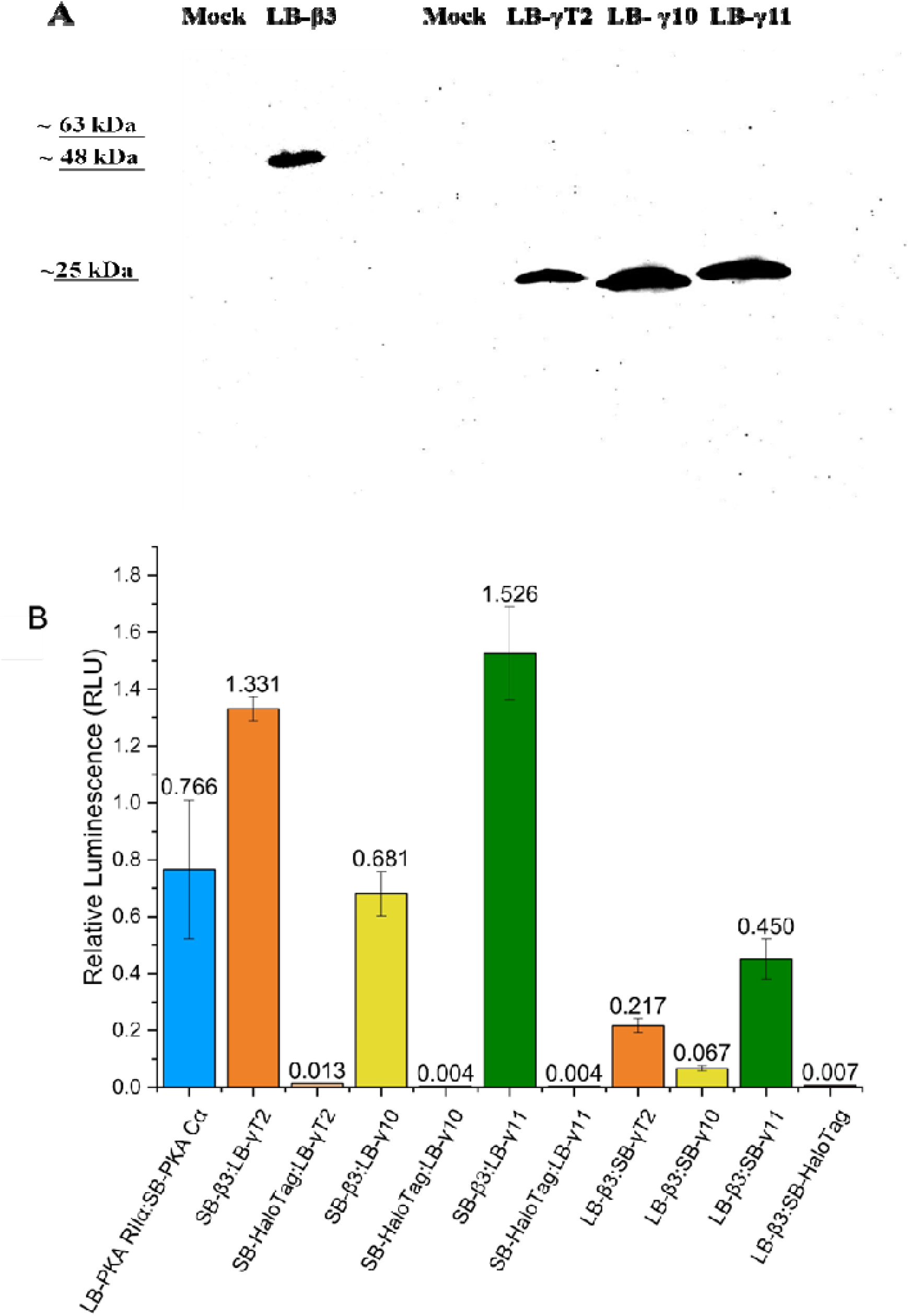
A: Expression of LB-β3 and LB-γ constructs in HEK cells detected by Western Blot. Anti-LB antibody was used with 1:2000 dilution on cell lysates. Non-transfected HEK cells (Mock) have been used as a negative control. Constructs shown to be expressed are detected at the expected size (∼55 kDa for the LB-β-subunit and ∼25 for the LB-γ subunits). B: NanoBiT βγ interaction assay. The results are shown as technical triplicates of relative luminescence units (RLU) for the tested NanoBiT constructs. Three biological replicates are shown in the supplement (Figure S7). LB-PKA RIIα:SB-PKA Cαis used as a positive control. As negative controls SB-HaloTag-LB-Protein fusion constructs were used.

**Table 2.**
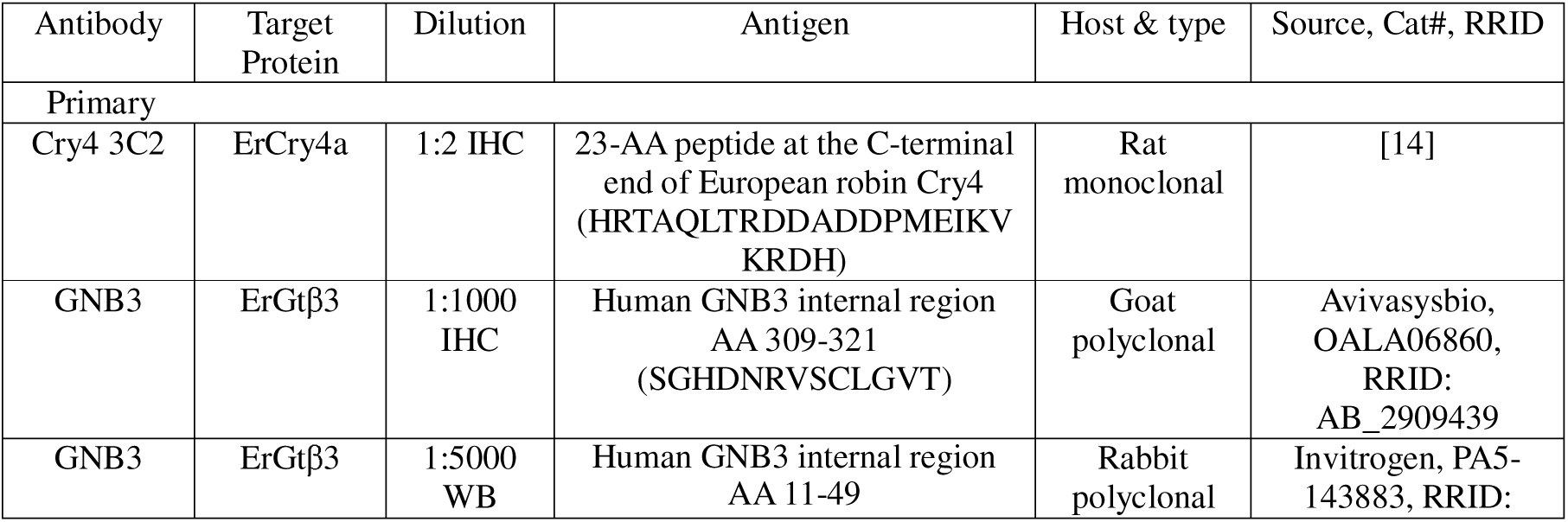
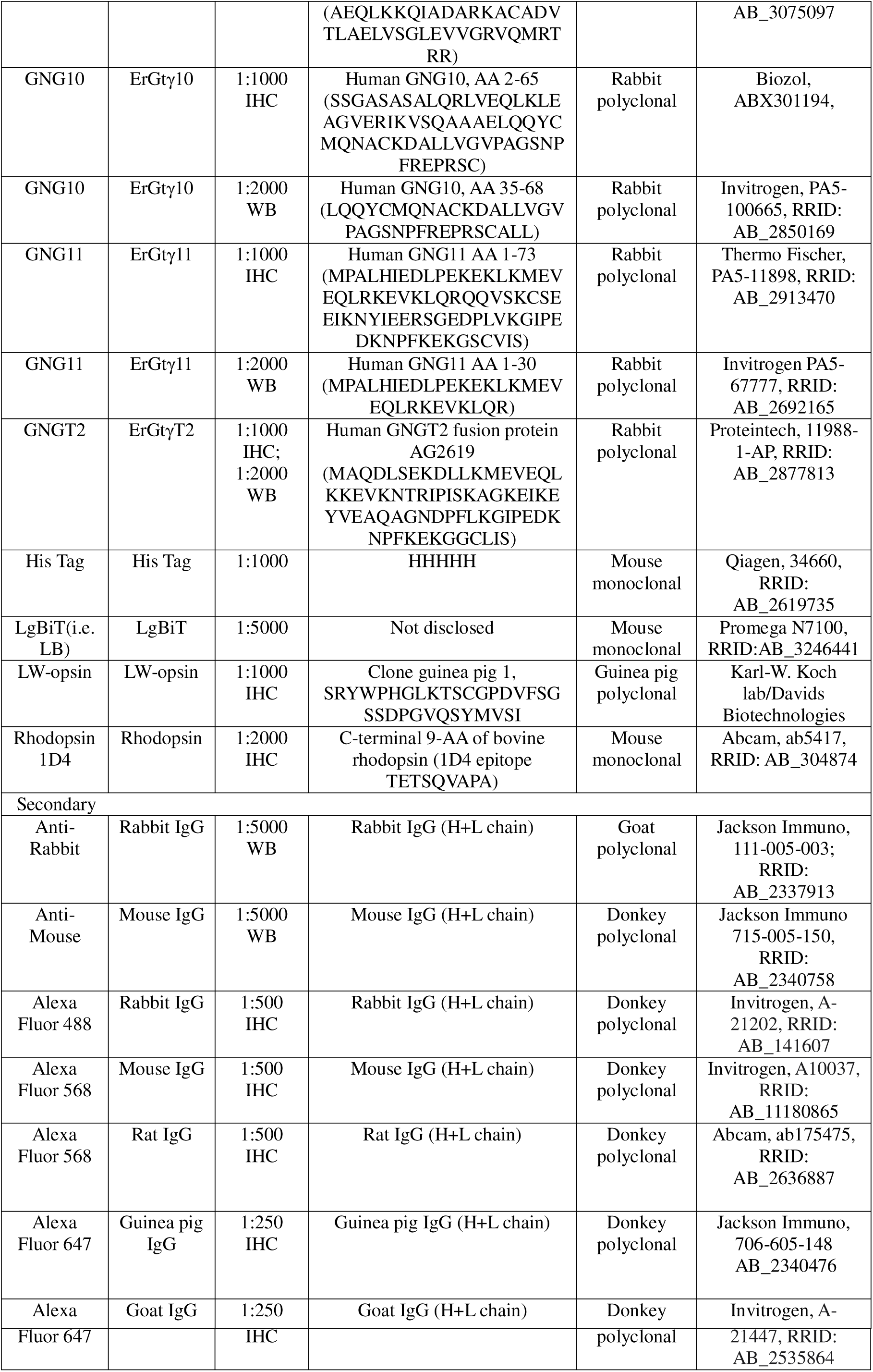
Antibodies used in the study. AA, amino acids; IHC – Used for Immunohistochemistry; WB - Used for Western Blot

The reason why only the N-terminus of the subunits has been tagged and tested is based on the known structure of Gtβγ-subunits, where the N-termini are in close contact [38]. Therefore, these constructs would provide an optimal distance for assembly of the NanoLuciferase fragments, when Gtβγ pairs have formed. Additionally, the CAAX motif in the Gtγ-subunits was preserved in the corresponding constructs. Therefore, the expressed Gtγ-subunits will be membrane-attached due to prenylation. In contrast, C-terminal-tagged Gtγ-subunits would be hindered in regard to prenylation, which is why C-terminus tagging was not considered as an option in this experimental approach [39].

## 3. Discussion

This is, to the best of our knowledge, the first study on the expression of G-protein subunits in any songbird species. Our scRNAseq results for migratory European robins show that *GNB3* is the main subunit predominantly expressed in cones, while *GNB1* was shown to be the main subunit expressed in the rods (Supplementary Figure 1). The expression of *GNB5* was detected in both rods and cones. This subunit, however, forms a complex with regulators of G protein signalling proteins (RGS) that control the GTPase activity of Gtα, which is important for the light-sensitivity of photoreceptor cells [25,40–42]. Regarding the Gtγ-subunits, our scRNAseq expression data solely show *GNG11* expression in rods, whereas *GNGT2* was dominant in cones. *GNG10* was neither expressed in rods nor cones, and overall, the expression was low in the retina. This is very much in line with an earlier study on the chicken retina, which showed *GNGT2* expressed in all photoreceptors, *GNG11* exclusively in rods and no photoreceptor expression of *GNG10* [29], suggesting that the expression of G-protein subunits in photoreceptors is largely conserved across different avian species.

Immunohistochemistry results supported the scRNAseq results. Antibodies directed against Gtβ3 and GtγT2 labelled rods and cones with strong staining in the outer segments. This is in line with an earlier study on the chicken retina which showed Gtβ3 expression in all photoreceptors [25]. As our scRNAseq data suggest minimal Gtβ3 expression in rods, this might represent a cross-reactivity of the anti-Gtβ3 with Gtβ1. The Gtγ11 antibody showed prominent staining exclusively in rods and only faintly labelled cones, yet we cannot exclude a possible cross-reactivity with other Gtγ-subunits. According to scRNAseq, transcripts of *GNGT2* and *GNG11* are both present in rods and cones, making a co-expression of both Gtγ-subunits likely. It is worth noting that Gtα2 and Cry4 co-localized in cone outer segments (Figure 2), in agreement with the Cry4 staining in Günther et al. (2018) [14], and supporting previous work showing a complex formation of Gtα2 and Cry4a on the protein level [34,43,44].

The anti-Gtγ10 antibody did not detect this isoform in any photoreceptor type of the European robin retina, and the transcript analysis showed its presence in retinal ganglion cells, oligodendrocytes and Müller cells, in line with earlier work in chicken [29]. Thus, our results do not support an involvement of Gtγ10 in the primary processes of vision, e.g. in the phototransduction cascade. It is also unlikely that Gtγ10 forms a complex with Cry4a in (double) cone photoreceptors and thereby participates in magnetoreception. However, the possibility of such an interaction was first suggested based on results from a yeast-two hybrid screening [19], which also identified Gtα2 as one of several candidate proteins capable of binding Cry4a. Considering that our results suggest absence of *GNG10* transcript in cones, the interaction of Gtγ10 with ErCry4 appears unlikely to occur in the retina of night-migratory European robins and the function of this isoform in the robin thus remains unknown. If, however, a Gtγ subunit were involved in magnetoreception, it would probably be GtγT2. A sequence alignment of GtγT2 and Gtγ10 (Supplementary Figure S5) showed that 42.6% amino acids are identical, with larger patches of higher similarity at the C-terminal half of the proteins, suggesting that what was identified as Gtγ10 (*GNG10*) in the yeast-two-hybrid screen might in fact be GtγT2.

Regarding subunit interactions, our *in vitro* studies showed that βγ-dimer formation occurred in living cells (Figure 6) but was not subunit-specific (Figures 5 and 6). Since our data suggest that all three tested Gtβ3/γ subunit combinations are possible, the formation of a functional heterodimer depends on the expression profiles in the specific cells, either in cone types or in rods. Therefore, our transcript analysis, immunohistochemistry and interaction studies are in agreement with a dimer pair of Gtβ3/GtγT2 in cones and Gtβ1/Gtγ11 and_/_or Gtβ1/GtγT1 in rods. These results agree in principle with the combinatorial diversity of βγ-interactions previously observed *in vitro* for human heterotrimeric G-proteins and investigated for formation of neurotensin receptor 1 G protein complexes [33].

In summary, we elucidate the diversity of heterotrimeric G-proteins in rods and cones of a night-migratory songbird and discard a functional role of Gtγ10 (encoded by *GNG10*) as a binding partner for European robin Cry4a.

## 4. Materials and Methods

### 4.1 Animals

Four adult European robins (*Erithacus rubecula)* were captured using mist nets in the vicinity of the University of Oldenburg under permit D7.2220/18 from the Lower Saxony State Department for Waterway, Coastal and Nature Conservation. Birds were housed in an indoor aviary under a natural light-dark cycle with *ad libitum* access to food and water. All birds were sacrificed by decapitation (two birds for transcriptomic analysis and two birds for immunohistochemistry). The tissue analysed in this study was obtained from birds that had been captured for other experimental purposes and subsequently became available for tissue collection. Thus, no randomization was performed, as birds were not experimentally assigned to groups. The sex and weight of all birds were recorded but were not experimental variables. All animal procedures complied with local, national and EU regulations for the use of animals in research and were approved by the Animal Care and Use Committees of the *Niedersächsisches Landesamt für Verbraucherschutz und Lebensmittelsicherheit (LAVES*, Oldenburg, Germany, 33.19-42502-04-22-00276). No animals were excluded or lost after inclusion in this study.

### 4.2 Single cell RNA sequencing of bird retina

#### 4.2.1 Tissue dissociation and single cell RNA Sequencing (scRNA)

The retinas were dissected immediately after decapitation, and the tissue was kept on ice. The retina tissue was dissociated into a single-cell suspension using a papain-based dissociation protocol with some slight modifications to account for varying cell counts following an adapted papain-based neural tissue dissociation protocol [45]. In brief, retinal tissue was dissociated enzymatically using papain in oxygenated Ringer’s solution (108 mM NaCl, 2.3 mM KCl, 1.8 mM MgSO_4_, 0.9 mM NaH_2_PO_4_, 25 mM NaHCO_3_, 30 mM glucose; adjusted to pH 7.4 with carbogen) supplemented with DNase I, followed by mechanical trituration and BSA-containing washes to obtain a nuclei suspension. Nuclei were isolated from the single cell suspension by following a standard detergent-based nuclei isolation protocol. Samples were homogenized in lysis buffer (10 mM Tris HCl, pH 7.4, 10 mM NaCl, 3 mM MgCl_2_, 0,1% Nonidet P40) using a Dounce homogenizer, diluted, incubated briefly with intermittent mixing, and filtered through a 70 µm strainer. The nuclei suspension was pelleted (500 g, 5 min, 4 °C), subjected to a second lysis step, and washed through sequential centrifugation steps using wash buffer (1 x PBS, 2% BSA, 0,2 U/µl RNase Inhibitor). The final nuclei suspension was resuspended, filtered through a 40 µm strainer, and collected in a fresh tube.

Single-nucleus 3′ gene expression libraries were prepared using the 10X Genomics Chromium Single Cell 3′ Gene Expression platform (Chromium Next GEM Single Cell 3 Reagent Kits v3.1 User guide CG000204). Libraries were sequenced on an Illumina platform as paired-end reads, with one read capturing the cell barcode and unique molecular identifier (UMI) and the other read covering the 3′ cDNA insert.

#### 4.2.2 Genome annotation

Functional gene annotation of the European robin reference genome [46] was performed using Maker version 3.1.4 (RRID:SCR_005309) [47]. This included soft masking the genome for transposable elements using RepeatMasker 4.1.2 (RRID:SCR_012954) with a library of transposable elements from the collared flycatcher [48] and the blue-capped cordon bleu [49]. For transcription level evidence we used ISOseq data from total retina, brain, muscle, lung, liver, heart and skin from one European robin, for which circular consensus sequences were generated to yield full-length transcripts [50]. Additionally, we included the transcriptome of chicken (*Gallus gallus*) as transcriptional evidence. All proteins in swissprot from birds were used as protein evidence. Using blast and exonerate through maker, these targets were mapped to the reference and used as input for the ab-initio gene predictors Augustus (RRID:SCR_008417 [51]) and snap (RRID:SCR_028812 [52]). Snap was additionally trained for two iterations, first using gene positions derived from evidence mappings and secondly using the output of the previous generation. The resulting genes were filtered for an annotation edit distance of less than 0.5, a metric provided by Maker to assess the goodness-of-fit of the annotation to the transcriptional evidence [47].

#### 4.2.3 Data integration and clustering

A count matrix was generated using the 10X Genomics CellRanger v7.2.0 software [53]. Using CellRanger mkref, the reference genome (Genebank accession GCA_903797595.1) and annotation was prepared for the fastq files to be aligned against. The output of filtered feature-barcode matrices was run through a standard Seurat pipeline [54] for quality control and downstream analysis. We defined thresholds for the feature and count values based on the distribution of the log_10_ transformed data such that we only included cells that had features in mean (log_10_nFeature) ± 2*SD (log_10_nFeature) and counts less than mean (log_10_nCount) + 2*SD (log_10_nCount). The data from both individuals were combined and integrated through Harmony [55]. Clustering was done using the Louvain algorithm with 0.5 as the resolution parameter to identify major clusters conservatively. Identities were assigned to each cluster based on the expression of canonical marker genes. For clusters that appeared to contain mixed cell types, individual cells were assigned identity based on marker genes.

#### 4.2.4 Data Analysis

We tested the expression of the following β- and γ-subunits of G proteins-*GNB1, GNB3, GNB4, GNB5, GNG2, GNG4, GNG7, GNG10, GNG11, GNG12* and *GNGT2* in different cell types in the samples. To assess cell type–specific expression of G protein subunits, we performed cluster-wise differential expression testing using a Wilcoxon rank-sum test, reporting log2 fold-change i.e. the logarithm of the ratio of expression levels between cell types and Benjamini–Hochberg–adjusted P values to identify subunits with significantly higher expression in a given cell type compared with all others.

### 4.3 Immunohistochemical stainings

#### 4.3.1 Tissue preparation and cryosectioning

Immediately after decapitation, eyes were enucleated and placed on filter paper (bottle-top filter, find listing). The anterior segment of the eye was removed with a razor blade, and the lens and vitreous humor were carefully separated from the eyecup. Eyecups were fixed in 4% paraformaldehyde (PFA) in phosphate-buffered saline (PBS, pH 7.4) for 30 min and washed thrice in PBS for 30 minutes each. For cryoprotection, eyecups were immersed subsequently in 10% and 20% sucrose in PBS until they sank and then in 30% sucrose overnight at 4 °C. Eyecups were stored in 30% sucrose in PBS at –20 °C until further use. For cryosectioning, eyecups were embedded in Tissue Tek O.C.T. Compound (Sakura Finetek Europe). Vertical retinal sections of 25 μm were cut on a cryostat and collected on Menzel SuperFrost Plus Adhesion slides (Thermo Fischer Scientific) and dried on a warming plate for 40-50 min. Slides were stored at –20 °C until further use.

#### 4.3.2 Immunohistochemistry

Retinal sections were washed in PBS 4-5 times for 10 min each. Primary antibodies (Table 2) were applied in PBS containing 0.3% Triton X-100 and incubated 1-3 days at 4°C. Following the primary antibody incubation, sections were washed 4-5 times for 10 min in PBS, followed by incubation with secondary antibodies diluted in PBS containing 0.3% Triton X-100 for 2 h at room temperature. After a final series of 4-5 times 10 min washing in PBS, slides were dipped in double distilled water for 1 min and mounted with Mowiol 8-88 (Sigma Aldrich) with DAPI (1.5 µg/ml) and coverslipped. Slides were stored at 4 °C until imaging. No specific staining was observed for the secondary antibody controls (Supplementary Figure S6).

#### 4.3.3 Confocal image acquisition and analysis

Images were acquired using the Leica TCS SP8 confocal laser scanning microscope equipped with an HC PL APO 63x/1.4 oil-immersion objective. Stacks were acquired with a zoom factor of 1.0 and an image resolution of 2048×2048 pixels. Z-step size was set to “system optimized”. Images are shown as average-intensity projections of confocal z-stacks. Image analysis was performed in Fiji (RRID:SCR_002285; [56]). Contrast was normalized using the *Enhance Contrast* function with the saturation threshold set to <1%.

### 4.4 Cloning of GNAT2, GNB3, GNG10, GNG11 and GNGT2

All used primers are available in Supplementary Materials (Table S3, Table S4). pFastBac™ HT B (Cat#10584027. Gibco, USA) vector was restricted with BamHI-HF and XhoI (Cat#R3136S and Cat#R0146S, New England Biolabs, USA) for 4 h at 37 °C. To amplify the coding sequences of *GNB3, GNG10, GNG11* and *GNGT2*, a PCR reaction was performed for 30 cycles. 10 ng of *E. robin* cDNA from retina was used as a template. Digested vectors were ligated with PCR products by Gibson assembly, and the samples were incubated for 2 h at 50 °C [57]. Assembled products were used to transform XL1 Blue Competent Cells (Cat#200236, Agilent Technologies, USA). Briefly, 20 µl of product was mixed with 5 µl of 10X CM Buffer (100 mM CaCl_2_, 40 mM MgCl_2_), filled up to 50 µl with ddH_2_O and incubated with 150 µl XL1 Blue competent cells for 30 min on ice. Cells were incubated for 1 min at 42 °C and, after heat shock, transferred to ice for an additional 10 min. After incubation, 800 µl of LB medium was added; cells were left on a shaker for an additional 30 min at 37 °C and plated on agar plates. The next day, colonies were picked, and Mini-Prep was performed (Cat# FG-90502, Nippon Genetics Europe, Germany). DNA was sent for sequencing, and positive clones were used for further experiments. A positive clone for GNB3 was used for deletion of the 6×His-Tag that was present in the pFastBac HT B vector. Deletion of the His-Tag was performed in a similar manner with appropriate primers, followed by a KLD reaction (Cat#M0554S, New England Biolabs, USA). The mixture was incubated for 30 min at 30 °C, and the success was confirmed by sequencing (Eurofins Scientific, Luxembourg). CAAX motif of γ-subunits was deleted in order to obtain soluble protein as described by [58]. For this, another PCR was performed, followed by KLD reaction and transformation was done in the same manner. Positive clones were used for further experiments.

For the NanoBit assay, PCR was performed to amplify the subunit coding sequence. Previously prepared pFastBac clones were used for the PCR reaction with the primers provided in Supplementary Table S5. MCS Starter System Vector templates [pBiT1.1-N (TK/LgBiT), pBiT2.1-N (TK/SmBiT), pBiT1.1-C (TK/LgBiT) and pBiT2.1-C (TK/SmBiT)] (Cat #N2014, Promega, USA) were digested with NheI and EcoRI (Cat#R3131S and Cat#R3101S, New England Biolabs, USA) at 37 °C for 2 h. The digested DNA was purified, and Gibson assembly was performed in order to ligate digested vectors with PCR products. The transformation of XL1 Blue cells was repeated as previously described. In total, 8 clones were prepared: N-term LgBitErGNB3, N-term SmBitErGNB3, N-term LgBitErGNGT2, N-term SmBitErGNGT2, N-term LgBitErGNG10, N-term SmBitErGNG10, N-term LgBitErGNG11, N-term SmBitErGNG11.

### 4.5 Expression of GNB3, GNG10, GNG11 and GNGT2 in ExpiSf9 cells

The method is based on a slight modification of the method by Iñiguez-Lluhi et al. (1992) [59]. Plasmids encoding the respective subunits were used to transform DH10Bac cells (Cat#10361012, Gibco, USA). Protein sequence information is available in Supplementary Table S2. 300 ng of DNA was used to transform 50 µl of thawed cells. The cells were incubated with the DNA for 30 min on ice. Afterwards, cells were incubated for 45 s at 42 °C and transferred on ice for an additional 2 min. 450 µl of Super Optimal Broth with Catabolite repression (S.O.C.) medium was added, and the mixture was incubated for 4 h at 37 °C and with medium agitation of 225 rpm. After the incubation, three 10-fold dilutions were prepared for spread plating on agar plates. 50 µg/ml kanamycin, 7 µg/ml gentamicin, 10 µg/ml tetracycline, 100 μg/ml 5-Bromo-3-indolyl-β-D-galactopyranosid (Bluo-Gal, Cat#15519028 Thermo Fischer, USA) and 40 µg/ml isopropyl-β-D-1-thiogalactopyranoside (IPTG) was pre-added to the agar plates. 100 µl of each culture was spread on plates. The plates were incubated for 3 days at 37 °C for blue-white selection. White colonies were picked to inoculate 3 ml LB medium with pre-added kanamycin, gentamicin and tetracycline and left to rotate overnight. The next day, Midi-prep was performed (Cat#19046, Qiagen, Germany). Concentrations of bacmid DNA obtained were between 2500-3500 ng/µl. For transfection, 2 ml of 1×10^6^ ExpiSf9 cells (Cat#A35243, Thermo Fisher Scientific, USA) were added to each well of a 6-well plate and left in the incubator to attach to the surface of the plate for 45 min at 27 °C. Meanwhile, a mixture of Cellfectin II (Cat#10362100, Thermo Fischer Scientific, USA) and bacmid DNA was prepared. 60 µl of Cellfectin II was mixed with 1 ml of ExpiSf™ CD medium (Cat#A3767801, Thermo Fischer Scientific, USA). 300 µl of the mixture was added to the other mixture containing 15 µl of bacmid DNA and 300 µl of medium and incubated for 40 min at RT. Additional 300 µl of medium was added afterwards. Medium from the wells was removed and 300 µl of the mixture was added to each well. Cells were incubated for 4 h at 27 °C. After incubation, the transfection mixture was removed, and 2 ml of fresh medium was added. Cells were incubated for 5 days, and P0 virus was harvested by centrifugation. P0 virus was used to infect 50 ml culture (1:10 dilution) containing 1.2×10^6^ cells, and the cells were left to shake for 3 days at 27 °C. P1 virus was collected and used to produce P2 virus in the same manner. For protein expression, 50 ml was infected with 0.5 ml of P2 virus (1:100) and incubated for 3 days. Cells were harvested; the pellet was washed with PBS (10 mM Na_3_PO_4_, 140 mM NaCl, 1 mM MgCl_2,_ pH = 7.4) and frozen at -80 °C until experiments.

### 4.6 Co-immunoprecipitation (Co-IP) studies

For cell lysis, the ExpiSf9 pellet with co-expressed subunits was resuspended in lysis buffer (20 mM Tris, 0.0025 mg/ml DNAse, 10 mM imidazole, 10 mM β-mercaptoethanol, 2.5 mM CHAPS, 300 mM NaCl pH = 7.5, Roche protease inhibitors). The cell pellet from 50 ml culture was resuspended in 1 ml of buffer and left to incubate on ice for 30 min. Cells were lysed by passing the cells through 25G needle for 5 min. Lysate was centrifuged at 100.000 g for 30 min, 4 °C. The supernatant was added to 50 μl of the pre-equilibrated Ni-Resin (Cat#88222, Thermo Fischer Scientific, USA). Resin pellet was washed with 3 x 500 μl of wash buffer (20 mM Tris, 300 mM NaCl, 50 mM imidazole, 10 mM β-mercaptoethanol, 2.5 mM CHAPS, pH = 7.5) and eluted with 50 μl of elution buffer (20 mM Tris, 300 mM NaCl, 300 mM imidazole, 10 mM BME, 2.5 mM CHAPS, pH = 7.5). Eluates were run on SDS-PAGE followed by Commassie staining and the rest was used for Co-IP.

Co-IP was performed using Invitrogen Dynabeads^TM^ Protein G (Cat#10003D, Thermo Fischer Scientific, USA). 10 µg of anti-His antibody (Cat#34660, Qiagen, Germany) was diluted in 200 µl TBS and incubated for 10 min at RT. Supernatant was removed and Dynabeads were resuspended in 200 µl of washing buffer (TBS with 0.02% Tween-20). Covalent crosslinking was performed. Dynabeads were washed 3 times with 500 µL 0.2 M sodium borate pH 9.2. Fresh solution of 20 mM DMP in 0.2 M sodium borate pH 9.2 was prepared. 1 ml of 20 mM DMP was added and incubated with rotation for 30 min at RT. Crosslinking was stopped with 2x500 µl washes of 0.2 M Tris-HCl pH 8.0. Dynabeads were additionally washed 3 times with 200 µl of washing buffer. Glycine pre-elution was used to remove non-crosslinked antibodies. Two washes with 500 µl 0.1 M glycine pH 2.0 were performed with subsequent three washes with 500 µl TBST to re-equilibrate. Eluate from preclearing was incubated with antibody-coupled Dynabeads for 1.5 h at 4 °C with rotation. Dynabeads were washed 5 times with IP buffer to remove unspecific proteins (20 mM Tris pH=7.5, 150 mM NaCl, 10 % glycerol, 2 mM EDTA 0.1% NP-40, Roche protease inhibitor). Then, the dynabeads were washed 5 times with wash buffer (20 mM Tris pH 7.5, 150 mM NaCl) to remove detergent, glycerol and EDTA. Beads were transferred to another tube and eluted with 50 µl of 2x SDS gel loading buffer. Eluates were used for Western blot.

### 4.7 NanoBit^®^ Protein-Protein Interaction Assay

NanoBit^®^ (NanoLuc^®^ Binary Technology) assay was performed to check interaction between G protein subunits [60]. Commercially available kit by Promega was used to produce LgBit and SmBit fused vectors of the subunits (in further text LB and SB stand for large <u>b</u>it and <u>s</u>mall <u>b</u>it, respectively). A positive control was provided consisting of the regulatory and catalytic domains of protein kinase A fused to the NanoBits resulting in the pair LB-PKA RIIα:SB-PKA Cα. As a negative control, HaloTag-SB plasmid was used to pair with the LB-subunit that was analysed in order to test for non-specific interactions. HEK293™ cells (Cat#R75007, Invitrogen, USA) were plated in a white 96-well plate (2.5×10^4^ cells per well, plated 24 h before transfection). Respective plasmids (1:1 ratio of LB- and SB-fused subunits) were first incubated with Lipofectamine™ 3000 transfection reagent (Cat#L3000001, Thermo Fischer Scientific, USA) for 20 minutes, prepared by mixing 0.2 µl P3000 reagent and 0.3 µl Lipofectamine per sample. After incubation, 10 µl of the mix was added to each well (corresponding to transfection of 100 ng of the pair per well). Cells were incubated for 24 h at 37 °C. The next day, Nano-Glo^®^ cell reagent (Cat#N2011, Promega, USA) was freshly prepared by diluting the Nano-Glo^®^ live cell substrate in Nano-Glo^®^ LCS dilution buffer (20-fold dilution). After addition of 25 µl of the reagent per well, the plate was incubated in the plate reader for 15 minutes at 37 °C for luminescence to develop. Luminescence was measured in Fluoroskan FL (Thermo Scientific, USA), using SkanIt Software 6.0.1., provided by the company. All experiments were performed in triplicates. Values were normalized to the mock control, containing only lipofectamine. Results were obtained in Relative Luminescence Units (RLU).

## Supplementary Materials

The following supplementary materials are available: Figure S1-UMAP plot showing the cell classes in the European robin retina using the cells from two individuals. Figure S2 - Violin plots showing the expression of G protein β-subunits in different cell classes. Y-axis: log-normalized transcript counts; Figure S3 - Violin plots showing the expression of G protein γ-subunits in different cell types. Y-axis: log-normalized transcript counts; Figure S4 – Immunohistochemical staining of central European robin retina for Gtγ10; Figure S5 – Sequence alignment of γT2 and γ10 subunits. Figure S6 – Immunohistochemical control labelling of central European robin retina using secondary antibodies only. Figure S7 – Biological replicates of NanoBit luminescence interaction studies. Table S1 – Markers for cell type classification in European Robin retina; Table S2 – G-Protein subunit sequences Table S3– Primer information for pFastBac cloning; Table S4 – Primer information for pFastBac CAAX motif deletion; Table S5 – Primer information for NanoBiT assay cloning;

## Data Availability Statement

The data referred to in this manuscript are present in the manuscript and in the supporting information. Unprocessed data are deposited on a server of the University of Oldenburg in accordance with the data policy of the Collaborative Research Centre SFB 1372 and are available on request from the corresponding author.

## Author Contributions

Conceptualization-ML, KD, KWK; formal analysis-SV, JJF, SK; supervision, ML, KD, KWK; investigation-SV, JJF, SK, GL, ÜGF, TB; methodology-SV, JJF, SK; visualization-SV, JJF, SK; writing, review and editing-SV, JJF, SK, HM, ML, KD, KWK. All authors have read and agreed to the published version of the manuscript.

## Acknowledgments

We would like to thank Dr. Rabea Bartölke for providing cDNA of *E. robin* and Alexander Rotov for his help with tissue collection for immunohistochemistry.

