## Supplementary information for "Expression patterns and interaction profiles of heterotrimeric transducin subunits in the retina of the European robin (*Erithacus rubecula*)"

**Figure S1**

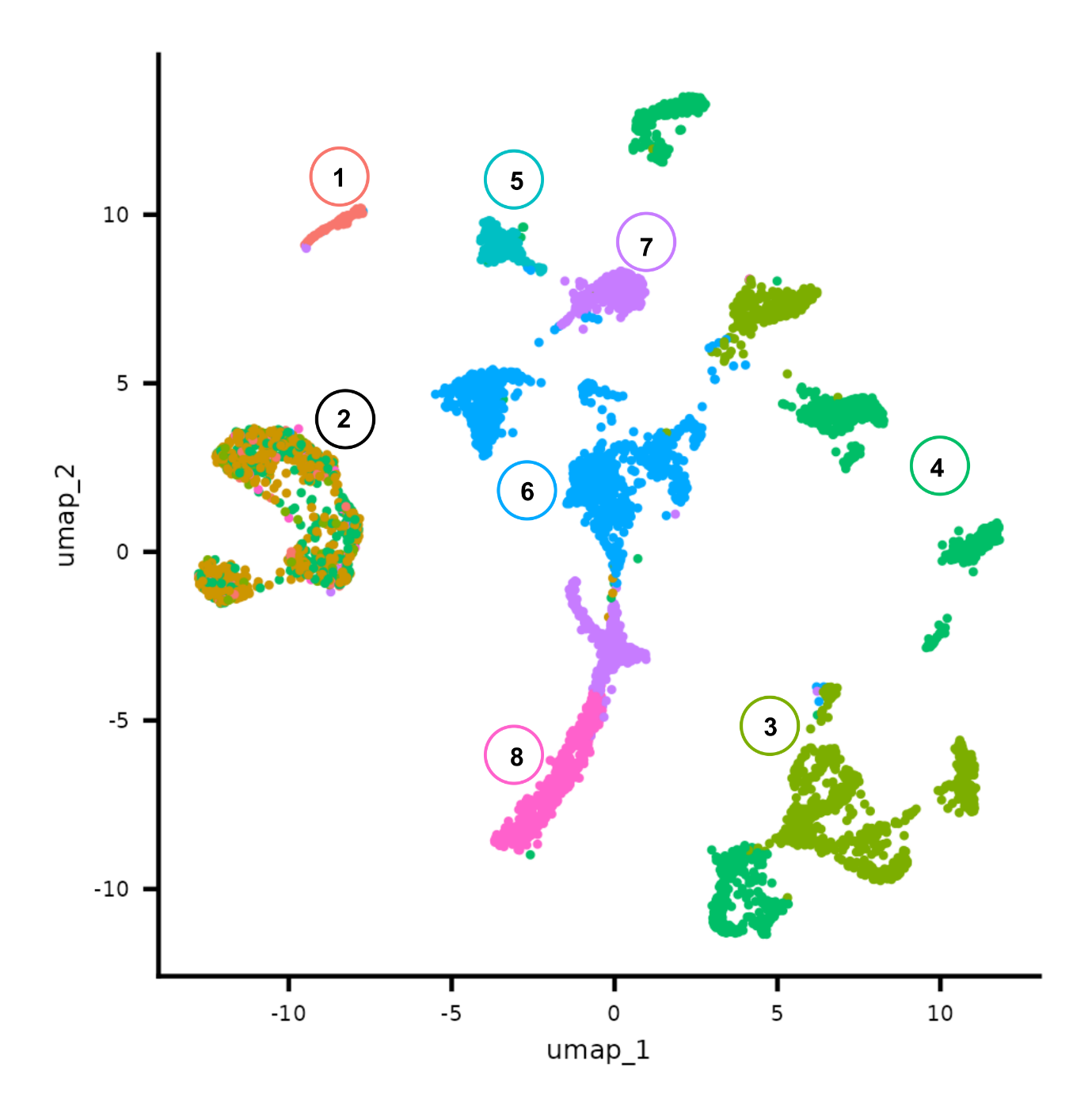
**Figure S1.** UMAP plot of scRNAseq expression data from 2 individual European robins characterising the different cell types of the European robin retina. (1) Rods. (2) Cones. (3) Bipolar Cells-Off. (4) Bipolar Cells-On. (5) Horizontal Cells. (6) Amacrine Cells. (7) Retinal Ganglion Cells. (8) Müller Cells.

**Figure S2**

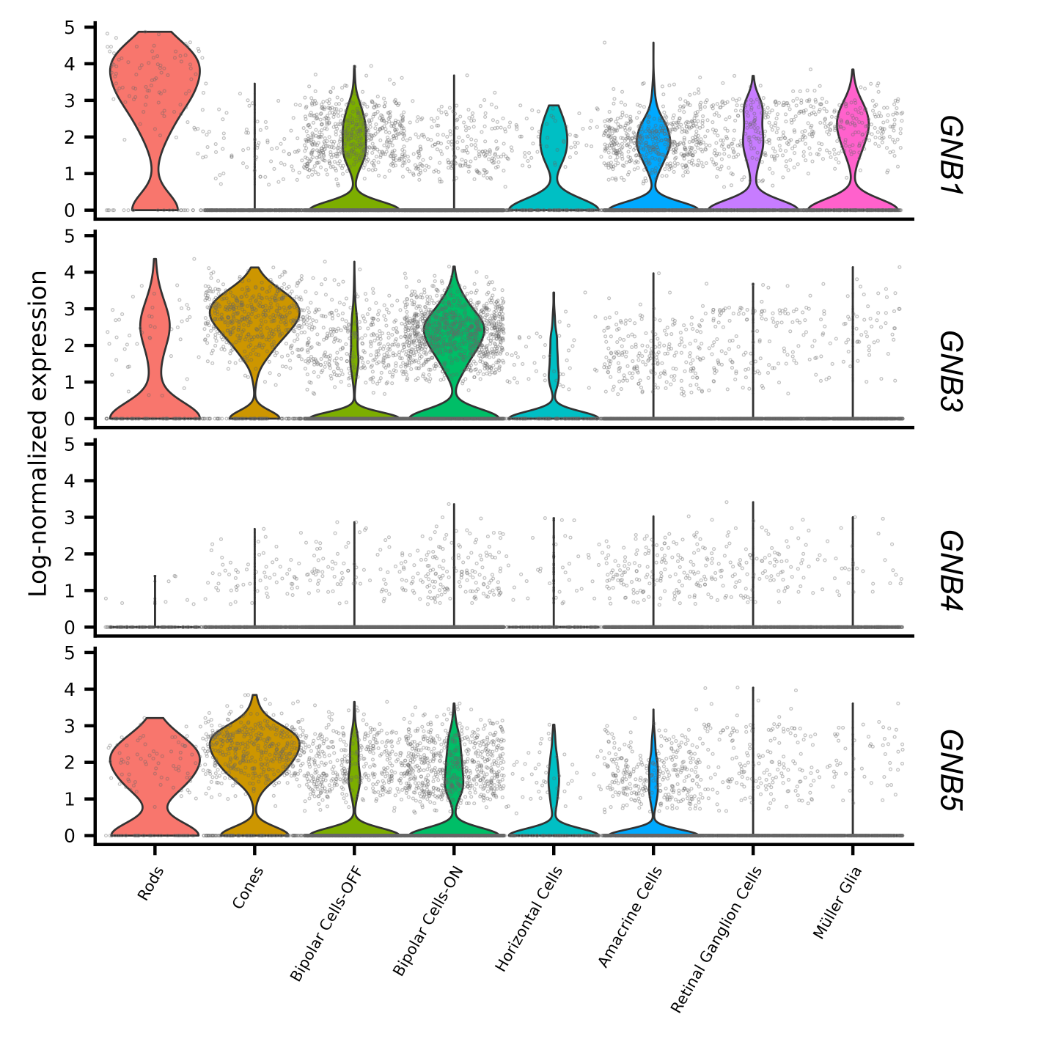

**Figure S2.** Violin plots showing the expression levels of G protein β-subunits in different cell types of European robin retina. Expression levels are plotted as log-normalized transcript counts.

**Figure S3**

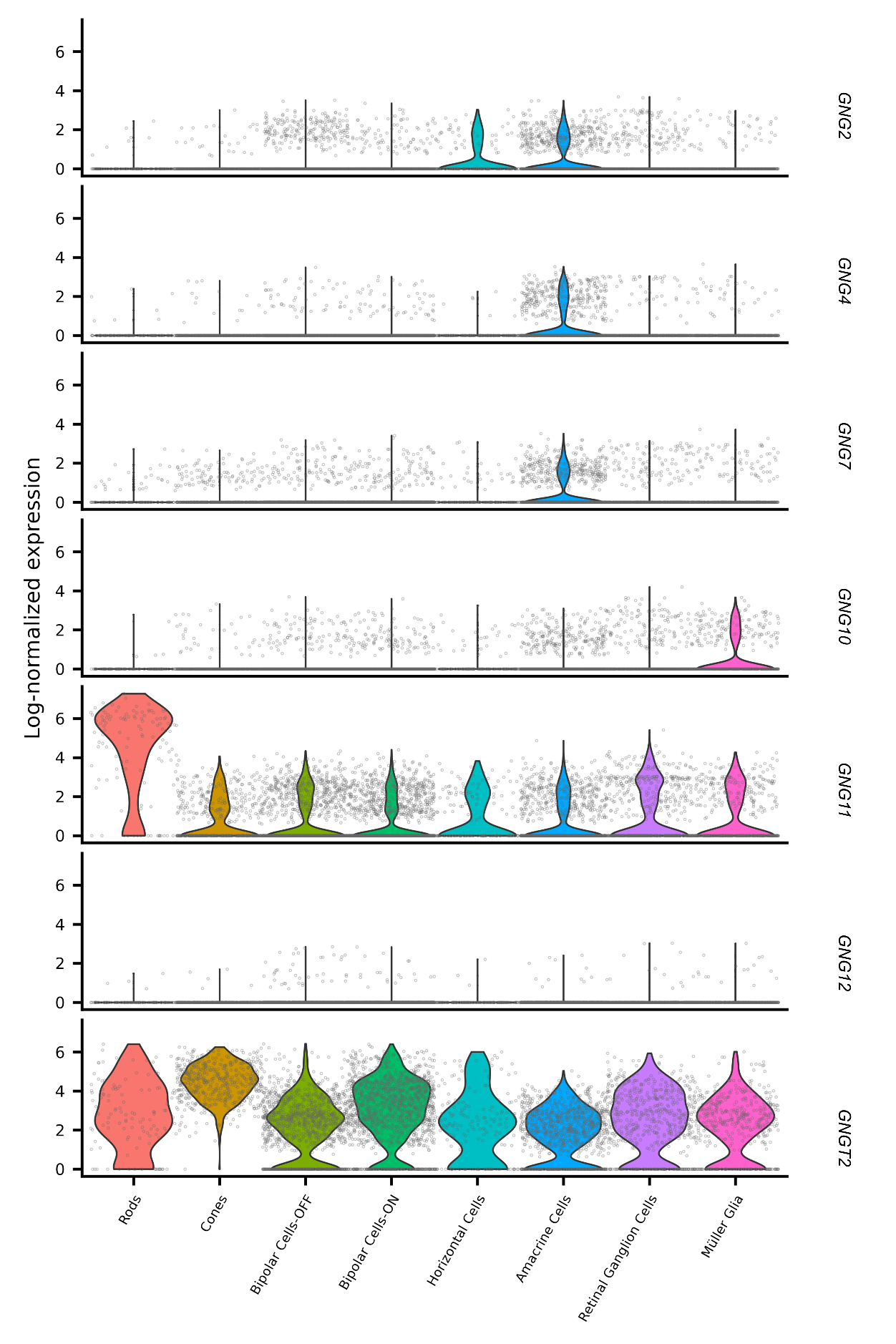

**Figure S3.** Violin plots showing the expression levels of G protein γ-subunits in different cell types of European robin retina. Expression levels are plotted as log-normalized transcript counts.

**Figure S4**

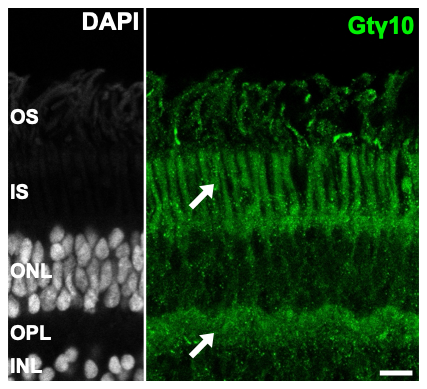

**Figure S4.** Immunohistochemical labelling of central European robin retina for Gtγ10. No specific labelling was detected, whereas diffuse signal was observed in the photoreceptor inner segments and endfeet (arrows). DAPI labelled sections (gray-scale) to visualize cell nuclei and retinal layers. OS, outer segments; IS, inner segments; ONL, outer nuclear layer; OPL, outer plexiform layer; INL, inner nuclear layer. Scale bar: 10 µm.

**Figure S5**

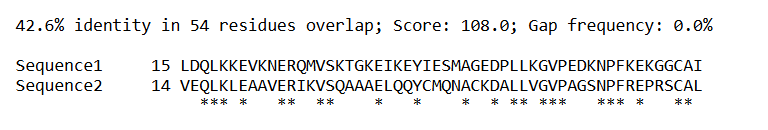

**Figure S5.** Sequence aligment of GtγT2 and Gtγ10 subunits. Aligment was done using Expasy (<https://web.expasy.org/sim/>)

**Figure S6**

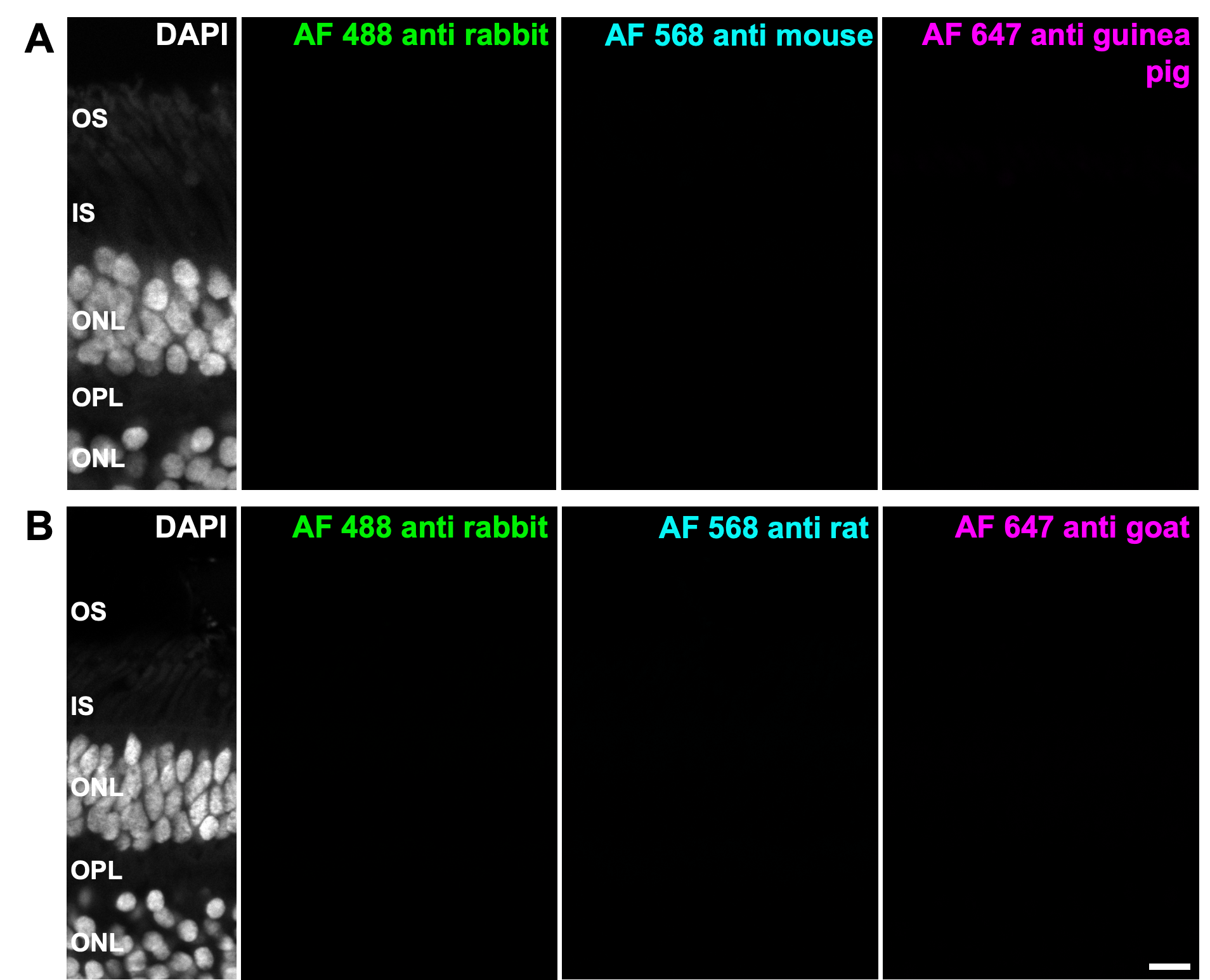

**Figure S6.** Immunohistochemical control labelling of central European robin retina using secondary antibodies only. (A) Retina incubated with Alexa 488 donkey anti-rabbit, Alexa 568 donkey anti-mouse and Alexa 647 donkey anti-guinea pig secondary antibodies. (B) Retina incubated with Alexa 488 donkey anti-mouse, Alexa 568 donkey anti-rat and Alexa 647 donkey anti-goat secondary antibodies. DAPI labelled sections (gray-scale) to visualize cell nuclei and retinal layers. Please note that control images were not contrast-enhanced. OS, outer segments; IS, inner segments; ONL, outer nuclear layer; OPL, outer plexiform layer; INL, inner nuclear layer. AF, Alexa Fluor. Scale bar: 10 µm.

**Figure S7**

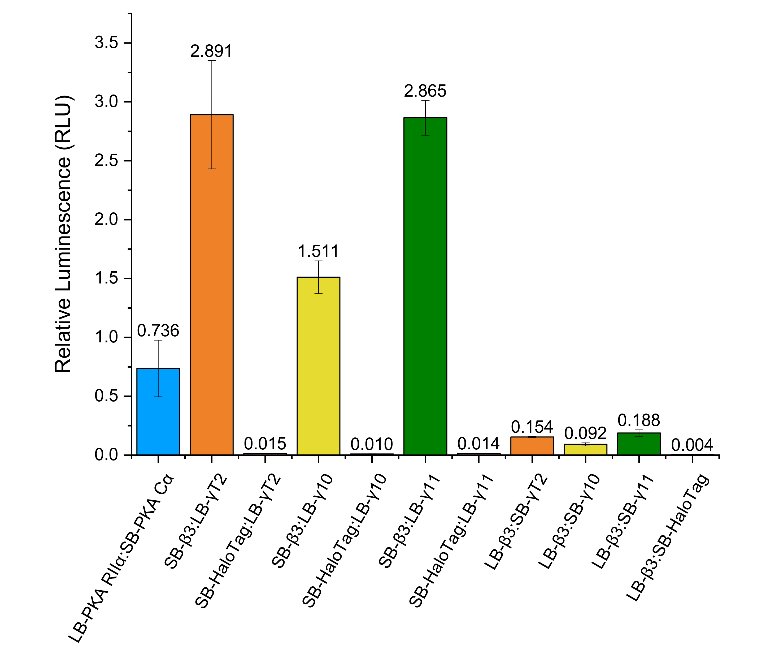

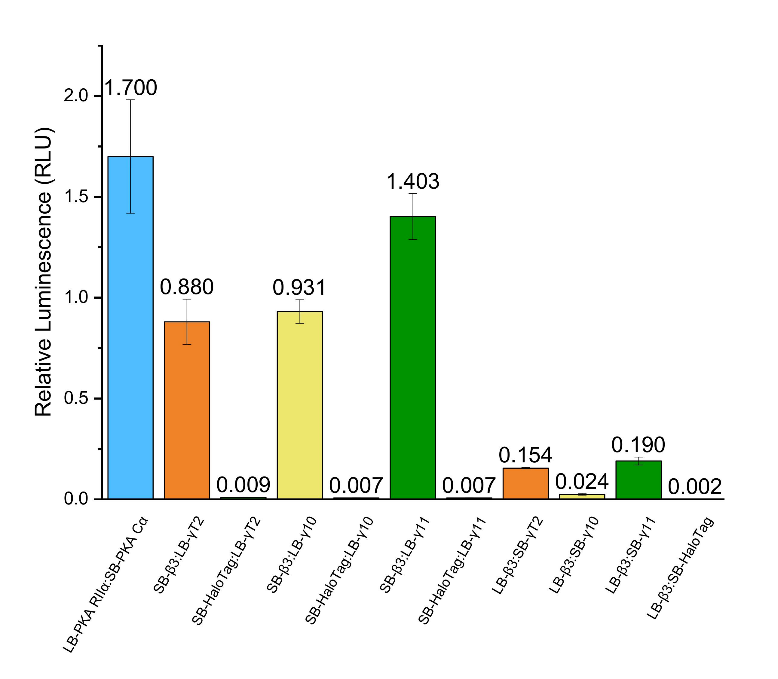

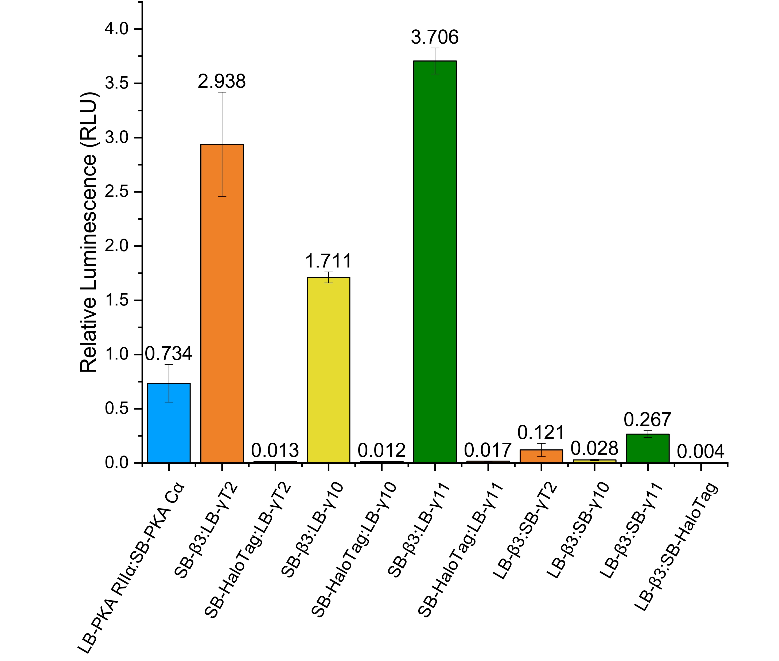

**Figure S7.** Three biological replicates of NanoBit luminescence interaction studies. Each set represents triplicates. Constructs and experimental setup were exactly as described in the main text.

**Table S1.** Markers for cell type classification in European Robin retina

| Cell_type | marker_name | marker_id (in annotation) |
| --- | --- | --- |
| Red SC | Red sensitive opsin | blast-LW-opsin |
| Red DC | MYLK | bEriRub00041045 |
| Blue SC | Blue Sensitive Opsin, SLC27A6 | OPN2SW; bEriRub00032869 |
| Green SC | PRA1 | bEriRub00040023 |
| UV SC | OPN1SW | bEriRub00011722 |
| Rod | RHO, PDE6B, PDE6G | bEriRub00027225; bEriRub00032820; bEriRub00035124 |
| Horizontal with axon | ONECUT3, LHX1 | bEriRub00041633; bEriRub00036556 |
| Horizontal without axon | ONECUT3, ISL1 | bEriRub00041633; bEriRub00021698 |
| Amacrine | SLC32A1 | bEriRub00031875 |
| Bipolar-ON | VSX2, OTX2, TRPM1 | bEriRub00015433; bEriRub00016246; bEriRub00028191 |
| Bipolar-OFF | VSX2, OTX2, GRIK1 | bEriRub00015433; bEriRub00016246; bEriRub00000036 |
| RGC | RBPMS, THY1 | bEriRub00013708; bEriRub00039134 |
| Oligodendrocytes | OLIG2 | bEriRub00003242 |
| Muller Glia | RLBP1 | bEriRub00027900 |

**Table S2.** G-Protein subunit sequences

| Protein Name | Accession Number (UniProt; European Nucleotide Archive) | Gene Name |
| --- | --- | --- |
| α isoform of transducin | A0A7K7GZ57_ERIRU; NWY74980 | *GNAT2* |
| β3 isoform of transducin | A0A7K7G5R6_ERIRU; NWY64895 | *GNB3* |
| γT2 isoform of transducin | To be added; OZ482522 | *GNGT2* |
| γ10 isoform of transducin | A0A7K7G7Z6_ERIRU; NWY65644 | *GNG10* |
| γ11 isoform of transducin | A0A7K7GRW7_ERIRU; NWY72457 | *GNG11* |

**Table S3.** Primer information for pFastBac cloning

| Primer Name | Primer Sequence |
| --- | --- |
| GNB3 (for) | 5‘-CAGGGCGCCATGGGATCCATGGGGGAAATGGAGCAGATG-3' |
| GNB3 (rev) | 5‘-GGTACCGCATGCCTCGAGTTAGTTCCAAATCTTGAGGAAACTG-3' |
| ΔHis GNB3 (for) | 5‘-ATGGGGGAAATGGAGCAGATG-3' |
| ΔHis GNB3 (rev) | 5‘-GGTTTCGGACCGAGATCCG-3' |
| GNGT2 (for) | 5‘-CAGGGCGCCATGGGATCCATGGCTCAGGACATGACAGAG-3' |
| GNGT2 (rev) | 5‘-GGTACCGCATGCCTCGAGTCAGCTGATGGCGCAGCCCC-3' |
| GNG10 (for) | 5‘-CAGGGCGCCATGGGATCCATGAGCTCGGGCGGCAGCC-3' |
| GNG10 (rev) | 5‘-GGTACCGCATGCCTCGAGTTAGAGCAGAGCACAGGATCGG-3' |
| GNG11 (for) | 5‘-CAGGGCGCCATGGGATCCATGCCAGCCATCAACATCGAGG-3' |
| GNG11 (rev) | 5‘-GGTACCGCATGCCTCGAGCTAAGCGATGACACAGCCTCC-3' |

**Table S4**. Primer information for pFastBac CAAX motif deletion

| Primer Name | Primer sequence |
| --- | --- |
| GNGT2, GNG10, GNG11 (for) | 5‘-TGACTCGAGGCATGCGGTACCAAG-3' |
| GNG10 (rev) | 5‘-GGATCGGGGTTCTCGGAAGGG-3' |
| GNG11 (rev) | 5‘-GCCTCCCTTCTCCTTGAACGG-3' |

**Table S5.** Primer information for NanoBiT assay cloning

| Primer Name | Primer Sequence |
| --- | --- |
| GNB3 (for) | 5‘-GAGCTCAGGGGAATTCAATGGGGGAAATGGAGCAGATG-3' |
| GNB3 (rev) | 5‘-CTAGAAGATCTGCTAGCTTAGTTCCAAATCTTGAGGAAAC-3' |
| GNGT2 (for) | 5‘-GAGCTCAGGGGAATTCAATGGCTCAGGACATGACAG-3' |
| GNGT2 (rev) | 5‘-CTAGAAGATCTGCTAGCTCAGCTGATGGCGCAGCCCC-3' |
| GNG10 (for) | 5‘-GAGCTCAGGGGAATTCAATGAGCTCGGGCGGCAGCC-3' |
| GNG10 (rev) | 5‘-CTAGAAGATCTGCTAGCTTAGAGCAGAGCACAGGATC-3' |
| GNG11 (for) | 5‘-GAGCTCAGGGGAATTCAATGCCATCAACATCG-3' |
| GNG11 (rev) | 5‘-CTAGAAGATCTGCTAGCCTAAGCGATGACACAGCC-3' |
